# A model of the entrance pupil of the human eye

**DOI:** 10.1101/325548

**Authors:** Geoffrey K Aguirre

## Abstract

The aperture stop of the iris is subject to refraction by the cornea, and thus an outside observer sees a virtual image: the “entrance pupil” of the eye. When viewed off-axis, the entrance pupil has an elliptical form. The precise appearance of the entrance pupil is a consequence of the anatomical and optical properties of the eye, and the relative positions of the eye and the observer. This paper presents a ray-traced model eye that provides the parameters of the entrance pupil ellipse for an observer at an arbitrary location. The model is able to reproduce empirical measurements of the shape of the entrance pupil with good accuracy. I demonstrate that accurate specification of the entrance pupil of a stationary eye requires modeling of corneal refraction, the misalignment of the visual and optical axes, and the non-circularity of the aperture stop. The model, including a three-dimensional ray-tracing function through quadric surfaces, is implemented in open-source MATLAB code.

## Introduction

When viewed from the outside, the aperture stop of the iris is seen through corneal refraction. The image of the stop is the entrance pupil of the eye. The precise appearance of the entrance pupil depends upon the anatomical and optical properties of the eye, as well as the relative positions of the eye and the observer. Characterization of the entrance pupil is relevant for modeling the off-axis image-forming properties of the eye (1), optimizing corneal surgery (2), and performing model-based eye-tracking (3).

The appearance of the entrance pupil as a function of horizontal viewing angle has been the subject of empirical measurement over some decades (4, 5). As an observer views the eye from an increasingly oblique angle, the pupil takes on an ever-more ellipitical appearance. Mathur and colleagues (6) measured the shape of the entrance pupil as a function of the nasal and temporal visual field positions of a camera moved about a stationary eye. The ratio of the minor to major axis length of the pupil ellipse (essentially the horizontal-to-vertical aspect ratio) was well-fit by a decentered and flattened cosine function of viewing angle. Fedtke and colleagues (7), using optical software, obtained a function with similar form by ray trace simulation of a rotationally symmetric cornea (8). As will be developed, the fit of this previous model to empirical data is imperfect.

Here I present a ray-traced model eye that provides the ellipse parameters of the entrance pupil as seen by an observer at an arbitrary location. The model accounts for individual variation in biometric parameters, including spherical refractive error. The location of the fovea is specified and used to derive the visual axis and line-of-sight of the eye. The open-source MATLAB code (https://github.com/gkaguirrelab/gkaModelEye) implements an invertible ray-tracing solution that can incorporate artificial lenses (e.g., spectacles, contacts) in the optical path between eye and observer. I simulate the circumstances of the Mathur and colleagues study (6) and show that their measurements can be replicated with good accuracy if the model incorporates *i)* corneal refraction, *ii*) the separation of the visual and optical axes, and *iii*) the non-circularity and tilt of the aperture stop.

## Model eye

In this section I describe the properties of a model eye specified in software. The parameters of the model are derived from prior reports (principally 9, 10), with some modification. Several aspects of the model vary with the spherical equivalent error of the eye. The coordinate system is centered on the optical axis, with the dimensions corresponding to axial, horizontal, and vertical position. Units are given in mm and degrees, with the exception of the tilt component of an ellipse, which is specified in radians. The quadric surfaces are specified in terms of the length of their radii (as opposed to the radius of curvature). Axial depth is given relative to the anterior-most point of the front surface of the cornea, which is assigned a depth position of zero. Parameters are presented for a right eye. Horizontal and vertical schematic sections of the model eye are presented in Figure 1.

**Figure 1.**
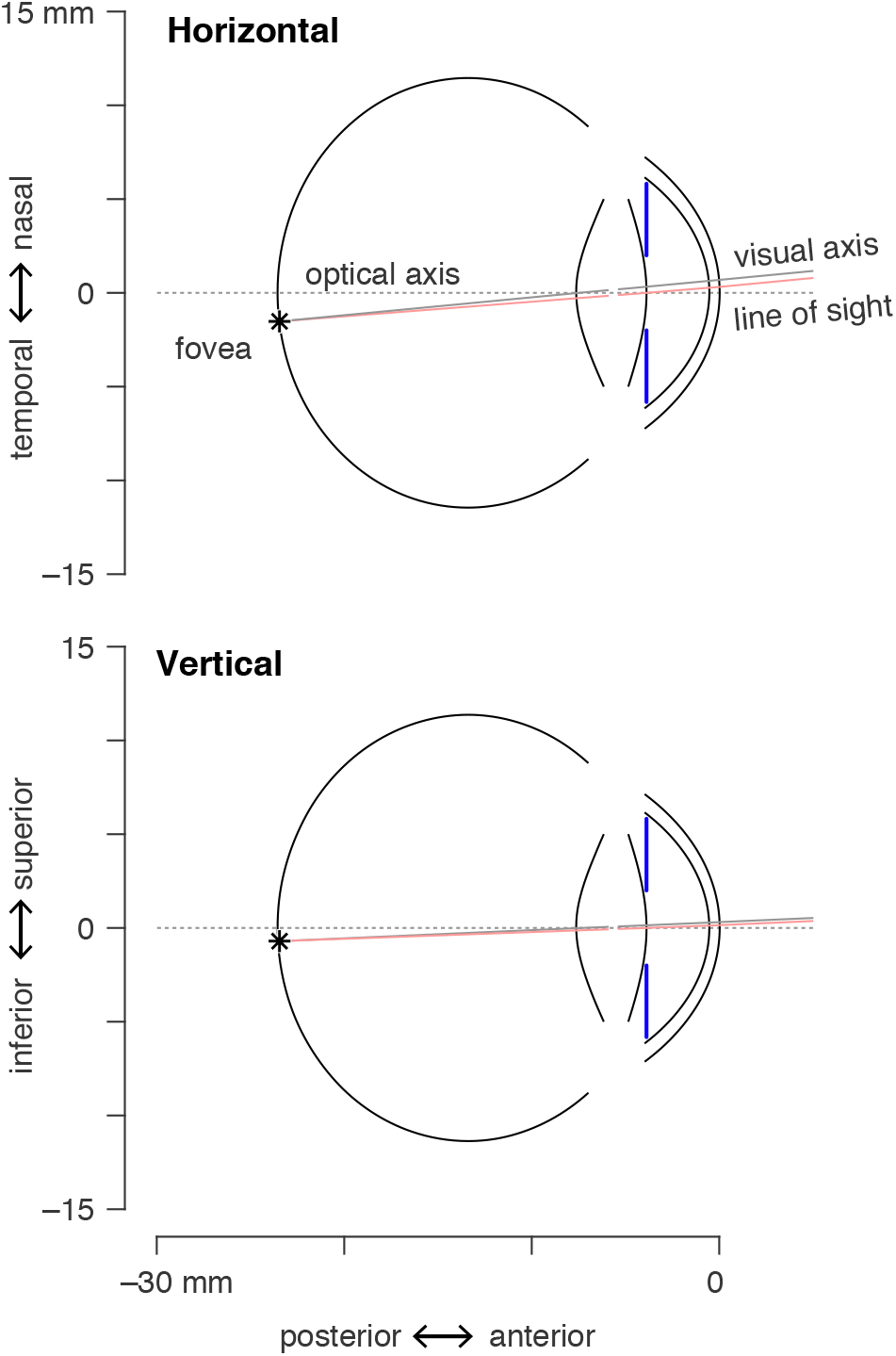
Schematic of the model eye. The components of the model are shown for an emmetropic right eye that is focused on a point 3 meters distant. The lens, cornea, vitreous chamber, and the aperture stop of the iris are aligned and centered on the optical axis (dotted line). The model specifies the location of the fovea (asterisk). Two rays are shown that connect the fovea to the fixation point. The visual axis (gray line) is a nodal ray of the optical system. The line of sight (pink line) passes through the center of the entrance pupil.

### Anatomical components

#### Cornea

The front surface of the cornea is modeled as a tri-axial ellipsoid, taking the “canonical representation” from Navarro and colleagues (9) for the emmetropic eye. Atchison (10) observed that the radius of curvature at the vertex of the cornea varies as a function of spherical ametropia, and provided the parameters for a rotationally-symmetric, prolate ellipsoid, with the radius of curvature (but not asphericity) varying linearly with the refractive error of the modeled eye. I calculated the proportional change in the Navarro values that would correspond to the effect of ametropia described by Atchison. This yields the following expression for the radii of the corneal front surface ellipsoid:

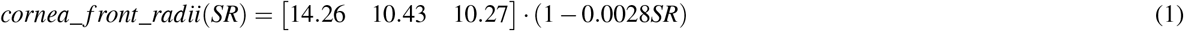

where *SR* is spherical refractive error in diopters, and the radii are given (here and subsequently) in the order of axial, horizontal, vertical. The apex of the corneal ellipsoid is on the optical axis.

Atchison (10) found that the back surface of the cornea did not vary by ametropia. Navarro and colleagues (9) do not provide posterior cornea parameters. Therefore, I scale the parameters provided by Atchison (10) to be proportional to the axial corneal radius specified by Navarro, yielding:

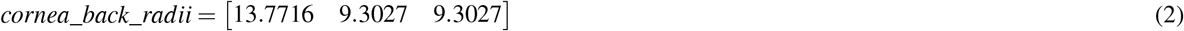

The center of the back cornea ellipsoid is positioned so that there is 0.55 mm of corneal thickness between the front and back surface of the cornea at the apex, following Atchison and colleagues (10).

#### Iris and aperture stop

The iris is modeled as a plane perpendicular to the optical axis (i.e., zero “iris angle”), positioned 3.9 mm posterior to the anterior surface of the cornea, at the location of the anterior surface of the lens for the cycloplegic eye (11).

The inner rim of the iris is the source of the image of the border of the entrance pupil. There is evidence that the entrance pupil is decentered nasally with respect to the optical axis, and that the center shifts temporally with dilation (reviewed in 12). The current model does not attempt to incorporate these measurements into the properties of the aperture stop. Instead, the aperture stop is modeled as centered and fixed on the optical axis.

The stop is, however, modeled as slightly non-circular. Wyatt (13) measured the average (across subject) ellipse parameters for the entrance pupil under dim and bright light conditions (with the visual axis of the eye aligned with camera axis). The pupil was found to be elliptical in both circumstances, with a horizontal long axis in bright light, and a vertical long axis in dim light. The elliptical eccentricity was less for the constricted as compared to the dilated pupil (non-linear eccentricity *ε* = 0.12 horizontal; 0.21 vertical; where the ratio of the minor to the major axis of an ellipse is equal to 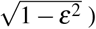. In the current model, I limit the eccentricity of the dilated entrance pupil to a slightly lower value (0.18) to provide a better fit to the measurements of Mathur and colleagues (6) (discussed below). Accounting for the refraction of the cornea (using ray-tracing simulations), I calculated the ellipse parameters (area, eccentricity, and tilt) for the aperture stop that would produce the entrance pupil appearance observed by Wyatt (13) at the two light levels. A hyperbolic tangent function of stop radius is then used to transition between the elliptical eccentricity of the constricted stop—through circular at an intermediate size—to the eccentricity of the dilated stop. Consistent with Wyatt (13), the tilt of the elliptical stop is horizontal (*θ* = 0) for all stop sizes below the circular intermediate point, and then vertical with a slight tilt of the top towards the nasal field for all larger stops.

The model therefore provides a stop that smoothly transitions in elliptical eccentricity and horizontal-to-vertical orientation (passing through circular) as the radius of the aperture varies from small to large following these functions:

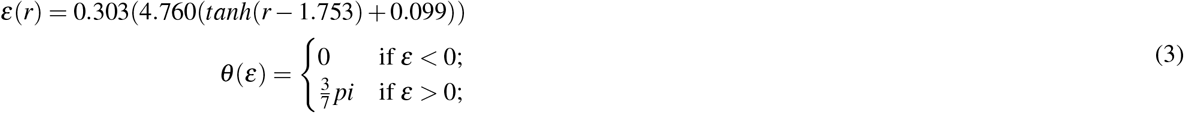

where *r* is the radius of the aperture stop in mm, *ε* is the non-linear eccentricity of the aperture ellipse, and *θ* is the tilt of the aperture ellipse (horizontal = 0; vertical = *pi*/2). The sign of *ε* is used to determine *θ*, although the absolute value of ε is used to generate the stop ellipse.

Although not required to determine the entrance pupil, the boundary of the visible iris is modeled for comparison with empirical images of the eye. The horizontal visible iris diameter (HVID) has been reported to be 11.8 mm (14). Bernd Bruckner of the company Appenzeller Kontaktlinsen AG measured HVID using photokeratoscopy in 461 people (15) and kindly supplied me with the individual subject measurements. These data are well fit with a Gaussian distribution and yield a mean visible iris radius of 5.91 mm and a standard deviation of 0.28 mm. After accounting for corneal magnification, the radius of the iris in the model is set to 5.57 mm.

#### Lens

The crystalline lens is modeled as a set of quadric surfaces. The anterior and posterior surfaces of the lens are modeled as one-half of a two-sheeted hyperboloid. The radii of these surfaces, their dependence upon age and the accommodative state of the eye, and thickness of the anterior and posterior portions of the lens, are taken from Navarro and colleagues (16). The interior of the lens is modeled with a gradient of refractive indices. This is realized as a set of ellipsoidal iso-indicial surfaces; 40 surfaces were used in the results presented here. The particular parameters are taken from Atchison (10), which were in turn derived from prior studies (17, 18). The lens is modeled as center-aligned with the optical axis. The axial position of the lens center is taken from Atchison (10).

The accommodative state of the model eye is set in terms of the distance of the point of best focus, established by adjusting the parameter “d” in the equations of Navarro and colleagues (16).

#### Vitreous chamber and retina surface

Atchison (10) provides the radii of curvature and asphericities of a biconic model of the vitreous chamber, which is de-centred and tilted with respect to the visual axis. With increasing negative refractive error (myopia), the vitreous chamber changes size, elongating in the axial direction and to a lesser extent in the vertical and horizontal directions. I model the vitreous chamber as an ellipsoid that is centered on and aligned with the optical axis, deriving the radii, and their dependence upon spherical refractive error, from equations 22-24 provided by Atchison (10).

#### Visual and line-of-sight axes

Our goal is to describe the appearance of the entrance pupil. Often, the pupil is observed while the subject under study has their eye directed at a fixation point. To model this, I define the location of the fovea on the retina and the visual and line-of-sight axes.

The visual axis is a nodal ray (19) of the eye that intersects the fovea. The angle α is the angle between the visual axis of the eye and the optical axis (12). The visual axis is generally displaced nasally and superiorly within the visual field relative to the optical axis. The horizontal displacement of the visual axis is 5 – 6° (20), vertical displacement is on the order of 2 – 3° (21). There is individual variation in these values.

Tabernero and colleagues (20) proposed that individual differences in the axial length of the eye are related to individual differences in α. Their model considers that, as the axial length of the eye increases, the axial distance of the fovea from the nodal points of the eye increases more than does the horizontal or vertical distance. This causes the angle between the visual and optical axes to grow smaller as the eye grows longer. In the current model, I adopt this theoretically motivated relationship with the form:

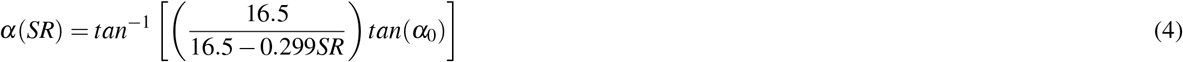

where *α*_0_ is the angle of the visual axis with respect to the optical axis in the emmetropic eye, and SR is spherical refractive error in diopters. The original Tabernero model is expressed in terms of axial length in millimeters; I converted this variable to units of diopters of spherical refractive error using the relationship given by Atchison (10), eq. 19. I set the *α*_0_ = 5.5° in the horizontal direction and *α*_0_ = 2.5° in the vertical direction. Ray tracing through the model eye is then used to assign the center of the fovea on the retinal surface as the destination of a ray that enters the cornea at the vertical and horizontal *α* angles.

When it is necessary to model the eye as focused upon a particular point in space, I obtain the line of sight for the model, which is the ray that connects the fovea and the fixation point via the entrance pupil

### Perspective projection and ray-tracing

Given the model eye, I then specify the appearance of the pupil in an image in terms of the parameters of an ellipse. This is accomplished by perspective projection of the eye to the sensor of a simulated camera, and then fitting an ellipse to points on the border of the pupil image. This projection requires us to specify the intrinsic properties of a camera and its position in space.

#### Perspective projection

A pinhole camera model is defined by an intrinsic matrix. This may be empirically measured for a particular camera using a resectioning approach (also termed “camera calibration”) (22, 23). If provided, the projection will model radial lens distortion (24), although the present simulations assume an ideal camera. The position of the camera relative to the eye is specified by a translation vector in world units, and the torsional rotation of the camera about its optical axis.

With these elements in place, the model defines points of equally spaced parametric angle on the boundary of the aperture stop. (These points are slightly unequally spaced in terms of their arc-length distance, but this is of no practical consequence.) The size of the aperture stop is specified as a radius in millimeters. As the stop is elliptical, radius is defined in terms of a circle of equivalent area. Five boundary points uniquely specify an ellipse and six or more are required to measure the deviation of the shape of the pupil boundary from an elliptical form. Sixteen points were used in this simulation, matching the number of points used by Mathur and colleagues to measure the entrance pupil ellipse (6). The boundary points are projected to the image plane and then fit with an ellipse (25), yielding the ellipse parameters.

#### Ray tracing

As viewed by the camera, the stop boundary points are subject to corneal refraction, and thus are virtual images. The cornea causes the imaged pupil to appear larger than its actual size, and displaces and distorts the shape of the pupil when the eye is viewed off-axis (3, 7). A similar and additional effect is introduced if the eye is observed through corrective lenses worn by the subject (e.g., contacts or spectacles).

To account for these effects, the projection incorporates an inverse ray-tracing solution. I model the cornea of the eye and any corrective lenses as quadric surfaces. The set of surfaces (with their radii, rotations, translations, and refractive indices) constitute the optical system. The routines provide refractive indices for the aqueous humor and cornea, as well as values for materials that are present in the optical path as artificial corrective lenses (e.g., hydrogel, polycarbonate, CR-39). The specific refractive index given the wavelength used to image the system is derived using the Cauchy equation, with parameters taken principally from Navarro and colleagues (16). A wavelength of 550 nm was used in the current simulations.

For the specified optical system, I implement 3D tracing for skew rays through a system of quadric surfaces (26). Given a boundary point on the stop, the angles with which the ray departs from the optical axis of the eye, and the spatial translation of the camera, we can compute how closely the ray passes the pinhole aperture of the camera (Figure 2). We search for the ray that exactly intersects the camera aperture and thus will uniquely be present in the resulting image (assuming a pinhole). The point of intersection of this ray at the last surface of the optical system is treated as the virtual image location of this point. In practice, the median ray trace solution finds a ray that passes within 0.0001 mm of the modeled camera pinhole. While greater precision could be obtained with longer computation time, this provides little practical benefit for model accuracy.

**Figure 2.**
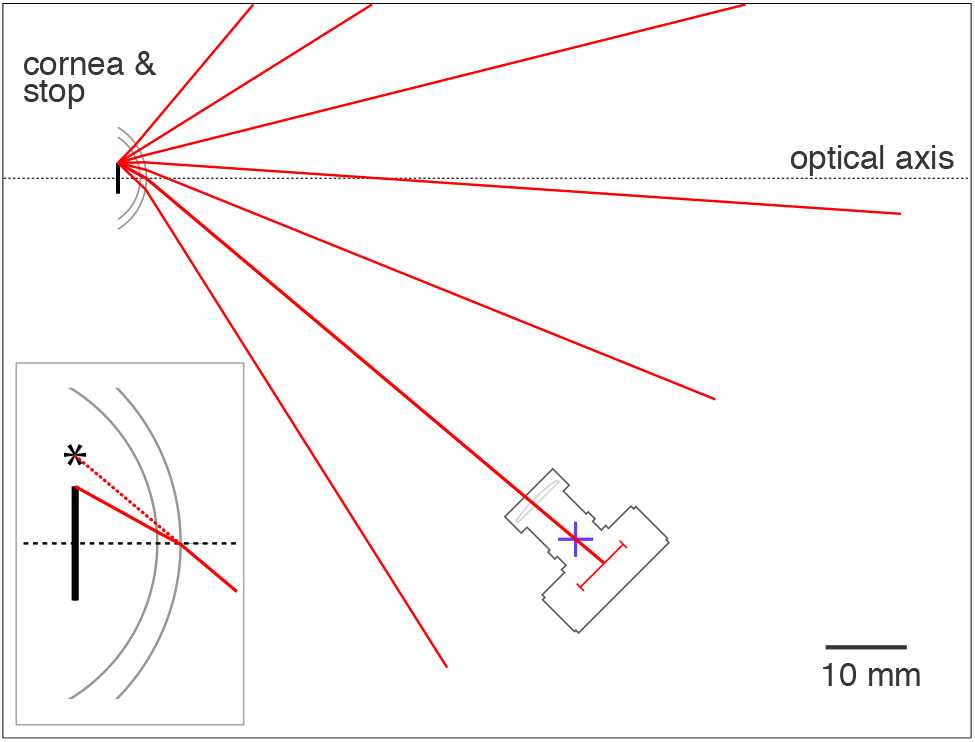
Calculation of the virtual image location of an aperture stop point. This 2D schematic shows the cornea and a 2 mm radius aperture stop of the model eye. A camera is positioned at a 45° viewing angle relative to the optical axis of the eye. The optical system is composed of the aqueous humor, the back and front surfaces of the cornea, and the air. We consider the set of rays that might originate from the edge of the aperture stop. Each of these rays depart from the stop border at some angle with respect to the optical axis of the eye. We can trace these rays through the optical system. Each ray will pass at a varying distance from the pinhole aperture of the camera (blue +). For a pinhole camera, only the ray that strikes the camera stop exactly (i.e., at a distance of zero) will contribute to the image. We therefore search across values of initial angle to find the ray that minimizes the intersection point distance. In this example system, a ray that departs the iris stop border at an angle of −30° with respect to the optical axis of the eye strikes the pin-hole of the camera after undergoing refraction by the cornea. *Inset bottom left.* The path of the ray after it exits the last surface of the optical system (in this case, the front surface of the cornea) creates a virtual image of the border point of the aperture stop that is displaced (asterisk) from the veridical location.

## Entrance pupil of a stationary eye

The ellipse parameters of the entrance pupil produced by the current model may be compared to empirical measurements. Mathur and colleagues (6) obtained images of the pupil of the right eye from 30 subjects. Each subject fixated a target 3 meters distant. A camera was positioned 100 mm away from the corneal apex and was moved to viewing angle positions in the temporal and nasal visual field within the horizontal plane. A viewing angle of zero was assigned to the position at which the line-of-sight of the subject was aligned with the optical axis of the camera. At each angular viewing position, the camera was translated slightly to be centered on the entrance pupil. An image of the pupil was obtained and then the border of the pupil marked by hand with 16 points, and these points fit with an ellipse. The pupil of the subject was pharmacologically dilated (and the eye rendered cycloplegic) with 1% cyclopentolate. The subjects had a range of spherical refractive errors, but the central tendency of the population was to slight myopia (−0.823 D). Figure 3A illustrates the geometry of the measurement.

**Figure 3.**
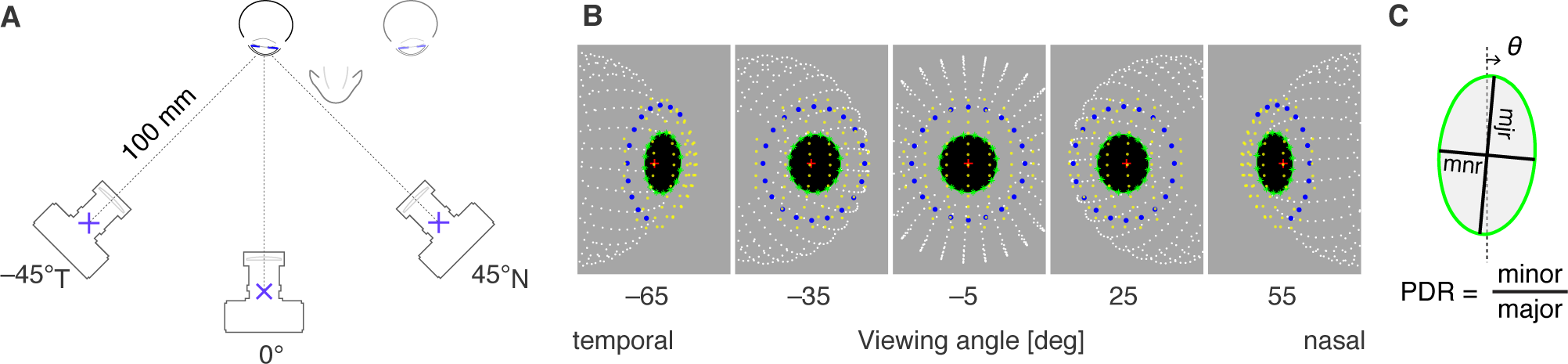
Simulation of measurements of Mathur and colleagues 2013. (A) A camera was positioned 100 mm from the cornea at different viewing angles within the temporal and nasal visual field. The line-of-sight of the eye was aligned with the optical axis of the camera at a viewing angle of zero. (B) The output of the current model is shown for equally spaced (30°) viewing angles from the temporal to nasal visual field. In the center image, viewed at −5°, the optical axes of the eye and camera are nearly aligned in the horizontal plane. The points shown correspond to the vitreous chamber (white), visible iris border (blue), entrance pupil border points and fitted ellipse (green), front corneal surface (yellow), and the center of the aperture stop (red plus). The iris border and pupil border points are virtual images, having been subjected to refraction from the cornea. Iris border points are missing in some views as these points encountered total internal reflection during ray tracing. (C) The pupil diameter ratio (PDR) is the ratio of the minor to the major axis of an ellipse fit to the entrance pupil. θ is the tilt of the major axis of the pupil ellipse. This value of *θ* is the empirically observed tilt of the entrance pupil ellipse, which is not to be confused with the specified value *θ* that is the modeled tilt of the elliptical aperture stop in Eq. 3.

The measurements of Mathur and colleagues (6) can be simulated in the current model. I created a right eye with a spherical refractive error of −0.823 D, and assumed an entrance pupil diameter of 6 mm to account for the effect of cyclopentolate dilation (27) (accounting for corneal magnification, this corresponds to a 5.3 mm diameter aperture stop). I set the accommodation of the eye such that the point of best focus was at 3 meters and rotated the eye so that the line-of-sight was aligned with the optical axis of the camera at a visual field position of zero. I then positioned the simulated camera as described in Mathur and colleagues (6). Figure 3B presents renderings of the model eye as viewed from the nasal and temporal visual field.

### Pupil diameter ratio

At each viewing angle Mathur and colleagues measured the ratio of the minor to the major axis of the entrance pupil, which they termed the pupil diameter ratio (PDR; Figure 3C). These measurements were combined across subjects and accurately fit by a parameterized cosine function. Figure 4A shows the resulting fit to the empirical data. The PDR decreases as a function of viewing angle, as the entrance pupil is tilted away from the camera and the horizontal axis is foreshortened. The observed entrance pupil follows a cosine function that is “both decentered by a few degrees and flatter by about 12% than the cosine of the viewing angle.” (6)

**Figure 4.**
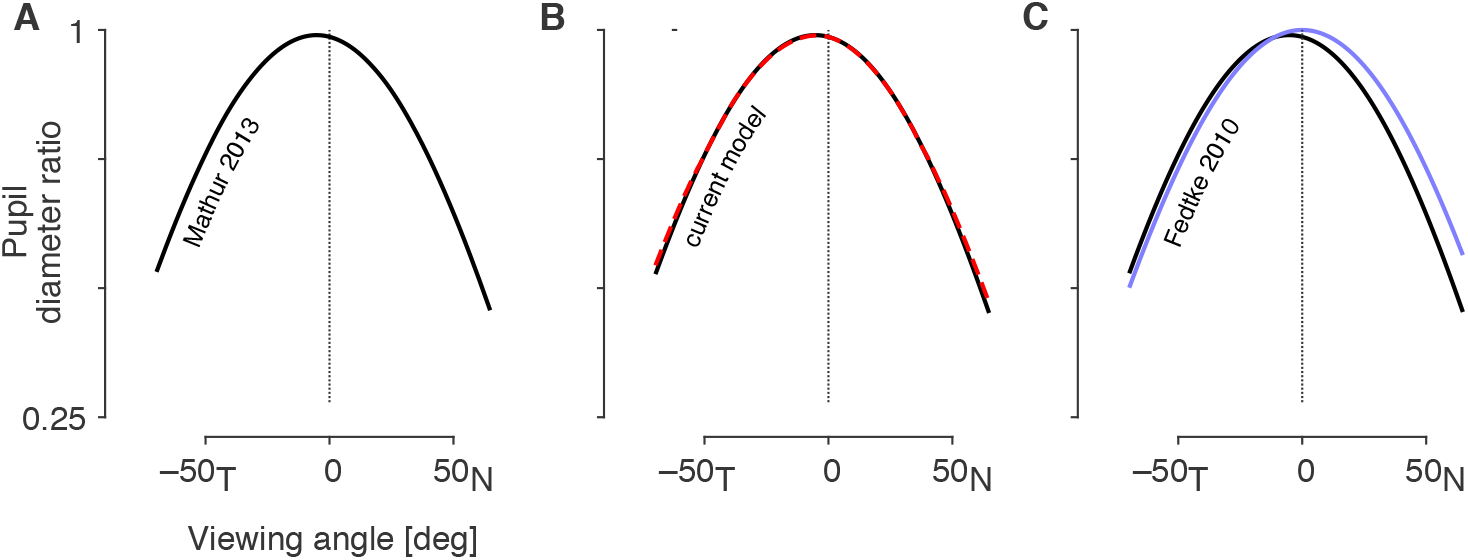
Empirical and modeled pupil diameter ratios. (A) The PDR as a function of viewing angle that was observed by Mathur and colleagues in 30 subjects. Note that the peak PDR is slightly less than one, indicating that the pupil always had an elliptical shape with a vertical major axis in these measurements. Also, the peak value is shifted away from a viewing angle of zero. (B) The current model provides the parameters of the entrance pupil ellipse. When the experimental conditions of Mathur and colleagues are simulated in the model, the resulting PDR function (dashed red line) agrees well with the empirical result (black). (C) In blue is the output of a model of the entrance pupil presented by Fedtke and colleagues in 2010.

Using the current model, I calculated the PDR for the entrance pupil with the camera positioned at a range of viewing angles. The model output (Figure 4B; red dashed line) agrees well with the empirical measurements. In prior work, Fedtke and colleagues (7) examined the properties of a simulated entrance pupil, assuming a rotationally symmetric cornea, a circular stop, and in which the visual and optical axes of the eye were equivalent. The PDR of their model as a function of viewing angle is plotted in blue in Figure 4C.

We may examine how components of the current model contribute to the replication of the empirical result. The simulation was repeated with successive addition of model components. The simplest model assumes a circular stop, no corneal refraction, and alignment of the optical axis and line-of-sight of the eye. Under these circumstances the entrance pupil behaves exactly as a function of the cosine of the viewing angle (Figure 5A). Next, we rotate the eye so that the line-of-sight is aligned with the optical axis of the camera when the viewing angle is zero (Figure 5B). Modeling of *α* (the misalignment of the visual and optical axes) decenters the PDR function. Addition of refraction by the cornea (Figure 5C) flattens the PDR function, as the entrance pupil now demonstrates magnification along the horizontal axis with greater viewing angles. Finally, the model is augmented with an elliptical, tilted stop (Figure 5D). Because the dilated stop is itself taller than it is wide, the PDR function is reduced overall, bringing the model into final alignment with the empirical result.

**Figure 5.**
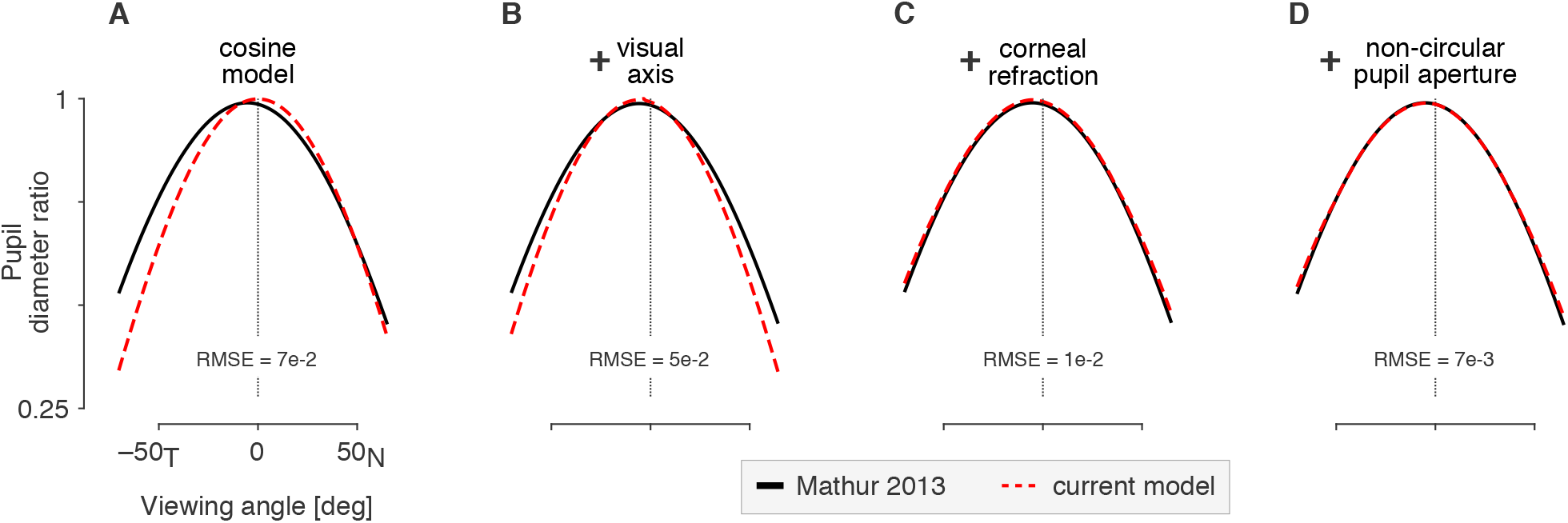
Effect of model components upon the entrance pupil diameter ratio function. In black is the Mathur and colleagues (2013) empirical result. The dashed red line is the behavior of the current model with the successive addition of model components (A-D). The root mean squared error (RMSE) between the model and the Mathur empirical result is given in each panel.

### Variation in the pupil diameter ratio function

Mathur and colleagues (6) found that the empirical PDR function could be fit with great accuracy by the expression:

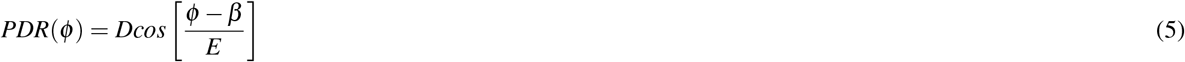

where *ϕ* is the viewing angle in degrees, *D* is the peak PDR, *β* is the viewing angle at which the peak PDR is observed (e.g., the “turning point” of the function), and 180E is the half-period of the fit in degrees. When fit to the output of the current model, we recover the parameters *D* = 0.99, *β* = −5.32, and *E* = 1.14, with an R^2^ = 0.99. These parameters are quite close to the those found (6) in the fit to empirical data (0.99, −5.30, and 1.12).

We may consider how biometric variation in the model changes the observed entrance pupil in terms of these parameters. The value *D* reflects the eccentricity and tilt of the stop ellipse (Eq. 3) and the vertical component of *α*. Wyatt (13) found individual variation in pupil shape, and by extension likely variation in the shape of the aperture stop in the iris. While variation in ellipticity of the stop will produce variations in *D*, overall this model component had a rather small effect upon the simulated PDR (Figure 5).

The value *β* is determined by the horizontal component of the misalignment of the visual and optical axes of the eye (angle *α*). This angle can vary substantially by subject and may have some relation to spherical refractive error. Figure 6A illustrates the PDR function associated with two different *β* values. Mathur and colleagues (6) obtained *β* for each of their 30 subjects and related this to subject variation in spherical refractive error. They found a negative correlation between these values, although this did not strongly account for the variance (R^2^ = 0.28). Figure 6B replots their data (gray points) and linear fit (black line). In the current model, the value assigned to *α* based upon spherical refractive error is made to follow the formula proposed by Tabernero and colleagues (20), producing the dependence of turning point upon spherical refractive error shown (red line). The current model also poorly accounts for individual variation in the value of *β* (R^2^ = 0.16).

**Figure 6.**
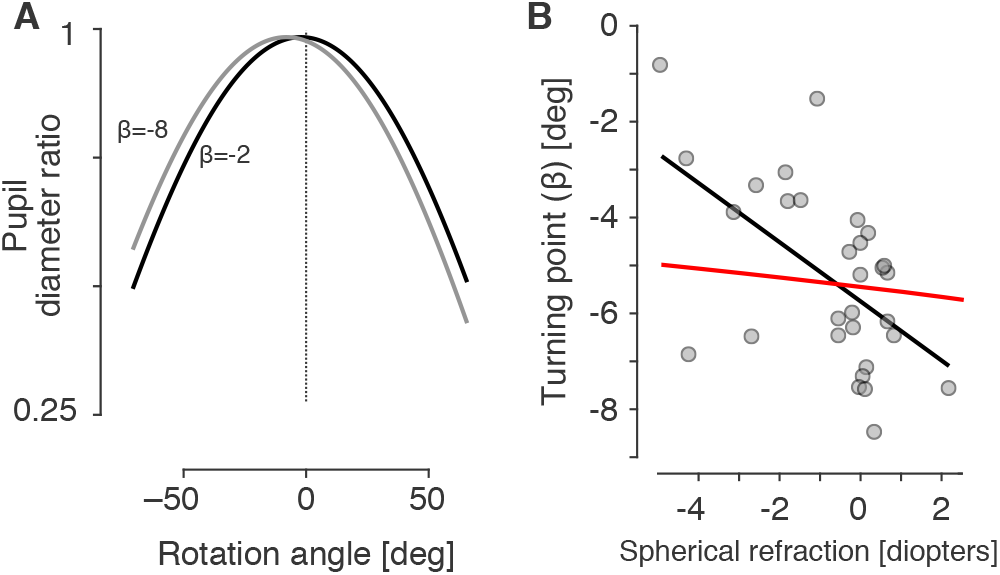
Variation in turning point. (A) The pupil diameter ratio was calculated using Equation 5 using *D* = 0.98, *E* = 1.14, and *β* values of −8 and −2. (B) In each of 30 subjects, Mathur and colleagues (2013) measured the viewing angle at which the peak pupil diameter ratio was observed (*β*), and related that value to the spherical refractive error of the subject. These data points (gray) are shown, along with the linear relationship between spherical error and turning point (blank line). Also shown are the turning point values returned by the current model when supplied with an eye of varying refractive error (red line).

The value *E* varies the “width” of the PDR function (Figure 7A). In their simulation, Fedtke and collagues (7) found that increasing the modeled aperture stop slightly widened the PDR function (i.e., the parameter *E* increased). I find the same effect in the current model (Figure 7B, top). Changes in the depth of the iris relative to the corneal surface had a minimal effect, suggesting that changes in accommodation (apart from a change in stop size) will have little effect upon the shape of the entrance pupil. I examined as well the effect of corneal radius as measured in the horizontal plane (Figure 7C, bottom). A modest change in the width of the PDR function seen for variation in the assumed corneal power. Given that the corneal surface is astigmatic, this indicates that there will be a small but measurable difference in the appearance of the entrance pupil along the vertical and horizontal dimensions.

**Figure 7.**
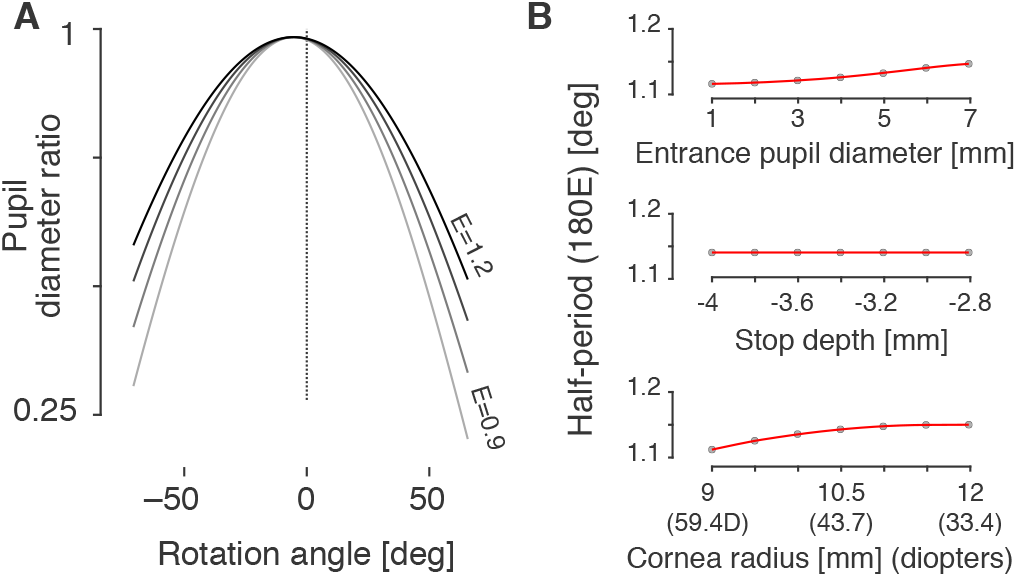
Variation in half-period width. (A) The pupil diameter ratio was calculated using Equation 5 using *D* = 0.98, *β* = −5.30, and *E* values ranging from 0.9 to 1.2. (B) The simulation was conducted with variation in pupil size, (*top*) iris depth (*middle*), and horizontal corneal radius (*bottom*). Variation in the *E* parameter assessed. Cubic-spline fits to the points are shown in red.

### Tilt of the entrance pupil ellipse

Mathur and colleagues (6) measured the “oblique component of pupil ellipticity” (termed component “C”), which reflects the tilt of the major axis of the pupil ellipse away from vertical. I calculated this component for the current model and found that the result compares quite well with the empirical measurements (Figure 8). This tilt component of the entrance pupil ellipse can be appreciated in the rendered eye depictions (Fig 3B), with the top of the pupil tilted towards the nasal field when the eye is viewed from the temporal field, and vice-a-versa. This tilt is almost entirely the result of modeling the vertical displacement of the visual axis from the optical axis; there is a small additional effect of the tilt of the dilated aperture stop.

**Figure 8:**
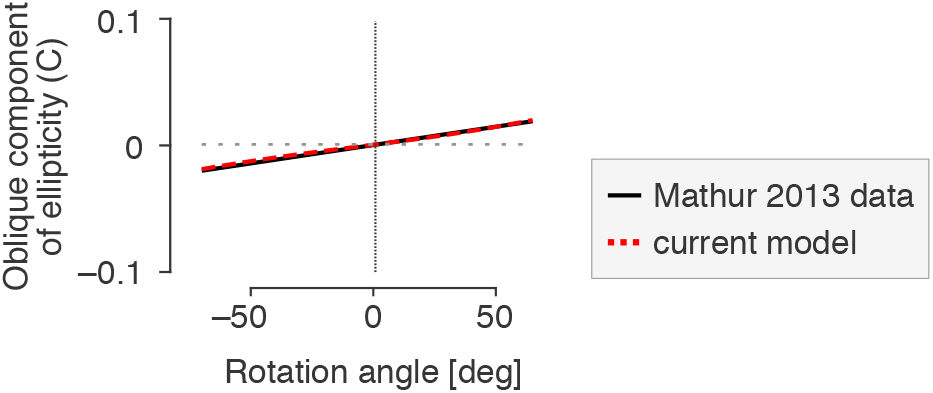
Oblique component of pupil ellipticity. The tilt of the pupil ellipse varies with viewing angle. Mathur and colleagues (2013) measured component “C” of the pupil ellipse, which was calculated as 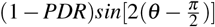, where *θ* is the tilt of the entrance pupil ellipse. Their observed C values are plotted in black. The current model (dashed red line) replicates the Mathur empirical result well (RMSE = 3e-3).

### Error in ellipse fitting to entrance pupil boundary

Finally, I examined how well the entrance pupil is described by an ellipse as a function of viewing angle by calculating the root mean squared distance of the entrance pupil border points from the fitted ellipse, expressed in millimeters (Figure 9). Overall, the ellipse fit error is small relative to the 6 mm diameter of the entrance pupil, although this error grows systematically as a function of viewing angle. Small discontinuities seen in the otherwise smooth function result from points at which one or more of the border points of the aperture stop encountered total internal reflection, leading to an ellipse fit performed upon fewer than 16 pupil perimeter points.

**Figure 9:**
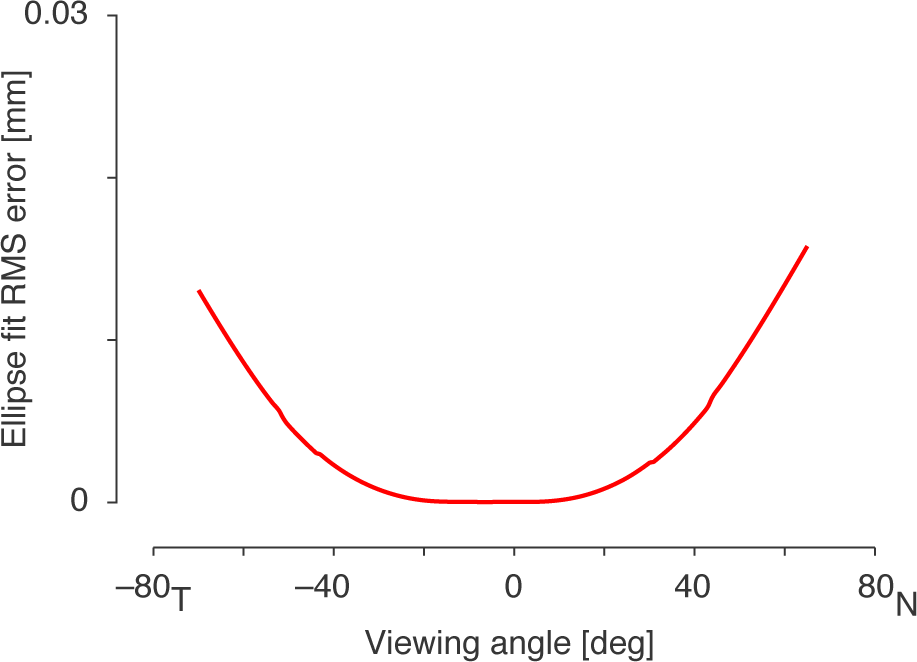
Ellipse fit error. The set of points that define the pupil boundary in the image are not exactly elliptical due to the refractive effects of the cornea. The root mean squared distance of the pupil boundary points from the best fitting ellipse as a function of viewing angle is shown. The fitting error is expressed in mm.

## Discussion

I find that a model eye is able to reproduce empirical measurements of the shape of the entrance pupil with good accuracy. More generally, the current study provides a forward model of the appearance of the pupil as observed by a camera at an arbitrary position.

Similar to the current study, Fedtke and colleagues (7) had the goal of using a ray-traced model eye to examine the properties of the entrance pupil as compared to empirical measures. The current study builds upon this prior work in a few respects. First, the anatomical accuracy of the anterior chamber of the model eye is advanced, incorporating an elliptical aperture stop and a tri-axial ellipsoid cornea. Second, the misalignment of the visual and optical axes is modeled. These refinements both improve the match to empirical observations (6) and support the generalization of the model to arbitrary pupil sizes and to viewing angles apart from the horizontal plane.

The model presented here provides the parameters of an ellipse that is fit to the border of the image of the entrance pupil. These parameters agree well with the empirical measurements of Mathur and colleagues (6). This does not require, however, that the true shape of the entrance pupil at high viewing angles is in fact elliptical. The measurements made in the Mathur study were obtained by fitting an ellipse to a set of 16 points that were manually defined around the pupil border. The quality of the fit was not reported. As both the current model and the Mathur measurements impose an ellipse fit, their agreement cannot confirm that the shape of the entrance pupil is in fact an ellipse. In the current model, an ellipse describes the border of the entrance pupil with very small fit error, even when the eye is viewed from 75 degrees in the periphery (Figure 9). The model may be inaccurate in this regard, however, as the peripheral cornea deviates from the ellipsoidal form used here (28). Indeed, at extreme angles, it is clear that the entrance pupil is decidedly non-elliptical in shape (4, 6). This is an area in which the current model could be improved.

While the goal in the current study was to account for the average appearance of the entrance pupil of the eye in a population, the model readily accommodates custom biometric measurements for any particular eye under study. Both the profile of the front surface of the cornea and the angle α have an effect upon the resulting entrance pupil appearance. Both have also been proposed to vary with spherical refractive error (10, 20), although this factor explains a relatively small amount of the individual difference. In the current work, I implemented the Tabernero and colleagues model of variation in α related to spherical refractive error (20). This model component did not perform well in describing the empirical variation in turning point observed by Mathur and colleagues (6). If α and spherical refractive error are in fact related, a larger set of empirical measurements, including cases of more extreme ametropia, will be needed to define the functional form.

The model is easily extended to account for rotation of the eye. There is an extensive literature in which anatomically-inspired models of the eye are used to inform gaze and pupil tracking systems (e.g., 3, 29). These applications have a particular focus upon determining the location of the first Purkinje image and the center of the entrance pupil, as these may be used to deduce gaze angle. While not studied here, these values are readily derived from the current model.

A feature of the current model is that it is implemented in open-source MATLAB code, as opposed to within proprietary optical software (e.g., Zemax). The ray-tracing operations of the model are conducted using compiled functions. As a result, the routines execute quickly (e.g., <10 msecs to compute the parameters of the entrance pupil ellipse on a standard laptop) and can thus form the basis of an “inverse model” that searches across values of eye rotation and stop radius to best fit an observed eye tracking image.

The biometric parameters that define the model were set almost entirely by reference to prior empirical measurements of the central tendency of studied populations. In an effort to best fit the pupil diameter ratio function of Mathur and colleagues (6), however, two model parameters were tuned “by hand”: the maximum, dilated entrance pupil ellipticity was held to a value slightly smaller than previously reported (13); and the value for vertical α was set to 2.5° in the emmetropic eye. Because these parameter values are biologically plausible and within the range of prior empirical measurements, I am optimistic that the model will generalize well to novel observations of entrance pupil appearance.

## Acknowledgements

Grant support provided by the NIH (U01EY025864, R01EY024681, P30EY001583).

## Author contributions statement

G.K.A. performed the study.

## Additional information

Competing interests The author declares no competing interests. Data availability The code used to perform these simulations and produce the figures may be found here: (https://github.com/gkaguirrelab/Aguirre_2019_NatSciReports)

